# Bio.Ontology - Python tools for enrichment analysis and visualization of ontologies

**DOI:** 10.1101/097139

**Authors:** Kamil Koziara, Julia Herman-Izycka, Bartek Wilczynski

## Abstract

**Motivation:** Functional annotation and enrichment analysis based on ontologies has become one of the standard methods of analysis of experimental results. Over the past decade, many methods have been proposed for statistical quantification of enrichment of different functional terms and many implementations of these methods are available. As the popularity of these methods grows, the need for tools facilitating their automation increases.

**Results:** We present a complete Python library for statistical enrichment analysis of gene sets and gene rankings compatible with most available biological ontologies. It allows the user to perform all necessary steps: reading the ontologies and gene annotations in multiple formats; performing enrichment analysis using various methods and visualizing the results as readable reports. Importantly, our library includes methods for correcting for multiple hypotheses testing including computation of False Discovery Rates.

**Availability:** The library is compatible with recent versions of python interpreter (≥ 2.6 or ≥ 3.3) and is available on github at: https://github.com/regulomics/biopython together with an API documentation and a tutorial. The sample galaxy installation can be found at http://regulomics.mimuw.edu.pl/wp/GO/.

## 1 Introduction

For the past several decades of explosive growth of molecular biology the problem of aggregating biological information and storing it in a usable format was a major challenge. Ontologies, i.e. graphical structures representing the hierarchy of terms in a research area together with a curated annotation of the connections between genes and ontology terms are growing and are becoming the *de facto* standard for knowledge representation for computer processing in the biomedical area. The publication of the original paper on Gene Ontology [ABB^+^00] marks an important step in the process of answering this need and providing the community with the tools for using it. Since that time, many scientists have engaged in multiple efforts to either create new ontologies or curate gene annotations to the correct terms of these ontologies most comprehensively exemplified by the OBO foundry repository [SAR^+^07].

Growing coverage of ontologies makes them more and more useful for automated analyses of gene sets and gene rankings. However, despite the quick increase of the number of proposed tools for ontology-based analyses the problem of finding a tool that can be easily integrated with other analyses remains unsolved for many researchers. In particular, many of the popular tools are either web-browser oriented like DAVID [HSL08] or desktop applications like GSEA [STM^+^05] or Ontologizer [BGVR08], making it the user responsibility to provide integration with the rest of the analysis pipeline. On the other end of the spectrum, there is a flurry of scattered libraries available online as a source code, often lacking support and stability required for trusting the results of the tool. In this work we present a new library for Ontology parsing and enrichment analysis that easily integrates with Biopython [CAC^+^09], one of the most popular python libraries for general bioinformatical analyses. It also includes an interface to the Galaxy web server [GNT10] allowing for integration with the users analyses both in the form of python scripts and galaxy workflows. In the following sections we will describe briefly the structure and functionality of the package together with a few examples of typical usage.

## 2 Implementation

The Bio.Ontology package consists of three main parts: input and output methods for reading the ontology and annotations in different formats; statistical methods for performing functional enrichment analyses on gene sets provided by the user and visualization methods for presenting the results. These parts revolve around the essential parts of workflow when carrying out the analyses (Fig. 1). Bio.Ontology allows user to find enriched terms in both: gene sets and gene rankings. To help with the problem of many genes having synonymous identifiers in different databases, our toolbox provides simple disambiguation method based on synonyms provided with the ontology annotation. Besides studied genes user needs an ontology and associations from genes to ontology terms. Presented software allows to use many different ontologies, many of which can be found at http://www.obofoundry.org. Bio.Ontology parses files in the most popular *obo* format and additionally it can handle the output generated by novel NeXO [DKS^+^12] software. Used ontology graph representation allows for editing: user can get subgraph of parsed graph induced either by specified set of terms or edges types. Additionally the graph can be visualized with the help of *graphviz* library. Bio.Ontology can read associations in standard *gaf* format. Depending on the size of annotation graph: smaller (e.g. single model organism) annotations can be stored in memory for fast access while larger annotation sets (e.g. Uniprot database) in a sqlite database to make the analysis feasible even on an average PC.

**Figure 1:**
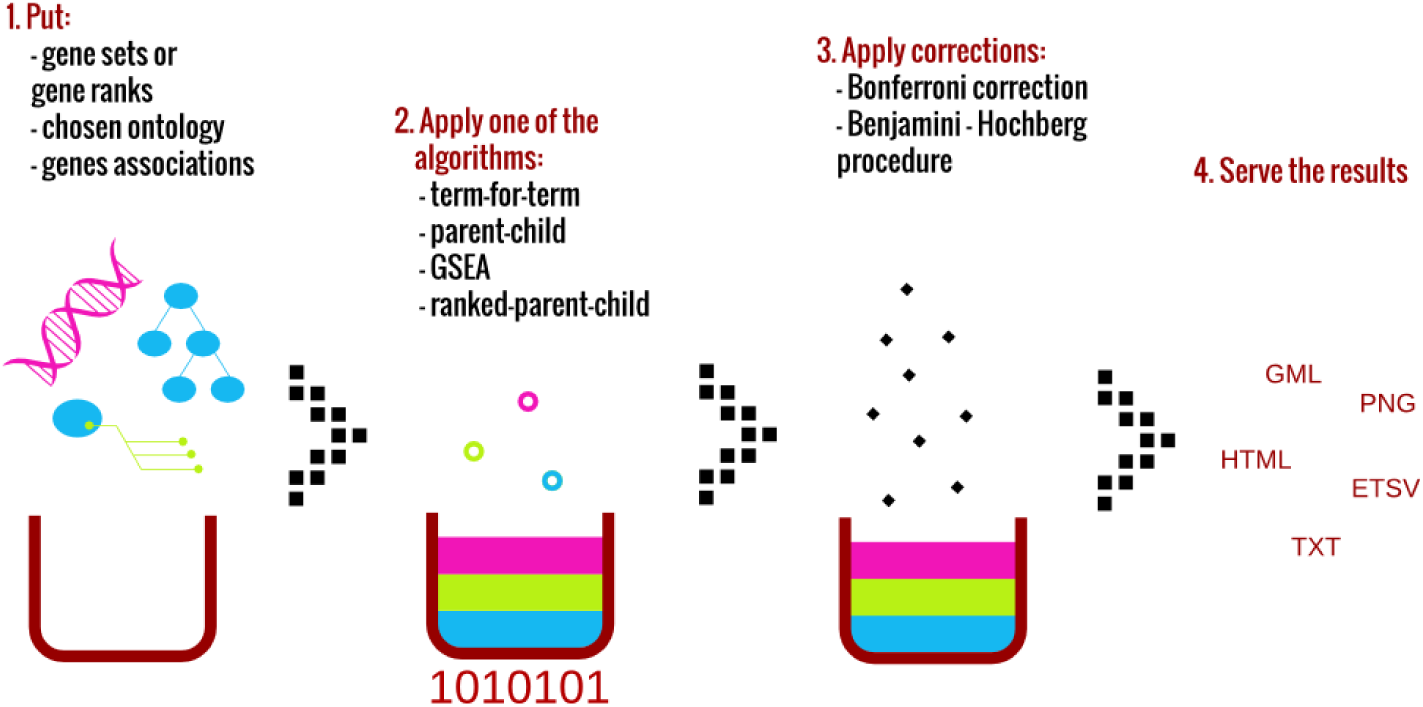
Bio.Ontology statistical term enrichment and visualization workflow.

After parsing the ontology and annotation data, Bio.Ontology allows the user to access the imported structures through python API using all biopython libraries. To facilitate the processing of ontologies, Bio.Ontology provides links to export the ontology into an object recognized by the networkx library [HSS08] allowing for wide array of graph algorithms and visualization methods.

As many users are interested in statistical enrichment analysis, Bio.Ontology provides built-in tools for various established methods for measuring statistical enrichment. This includes two methods for gene sets analysis: standard and most widely adopted *term-for-term* method based on one-tailed Fisher’s exact test and a more advanced *parent-child* [GBRV07] method which takes the hierarchical dependencies between the ontology terms into account when calculating the significance of enrichment. As it has become a standard practice in recent years, Bio.Ontology also allows for gene rankings analysis. Two methods available are: *GSEA* method [STM^+^05] and novel *ranked-parent-child* method developed with the toolbox. All of these are described in more detail in methods section.

Functional enrichment analysis usually involves testing large number of statistical hypotheses, standard corrections for multiple hypothesis testing are provided in our implementation. User can choose between Bonferroni correction and Benjamini-Hochberg procedure for controlling false discovery rate (FDR) [BH95]. Additionally, the *GSEA* method is coupled with its own method for controlling FDR based on resampling statistics.

All of the features mentioned so far can be also utilized from the provided interface to the Galaxy server [GNT10]. We provide a demo installation of the enrichment analysis tools on our website^1^, however galaxy users deploying their own servers can also install our package on their servers.

Obtained results can be filtered by p-value or FDR and then visualized. Bio.Ontology can generate reports as plain text or as *html* pages. Another way is to visualize enriched terms as graphs using either *graphviz* (an example in Fig.2 to put the results straight into image file or export them into *gml* format which can be later imported and edited using tools as *Cytoscape* [SMO^+^03]. Generated results can be as well exported to *etsv* (extended tab separated values) files and then imported again in the future.

**Figure 2:**
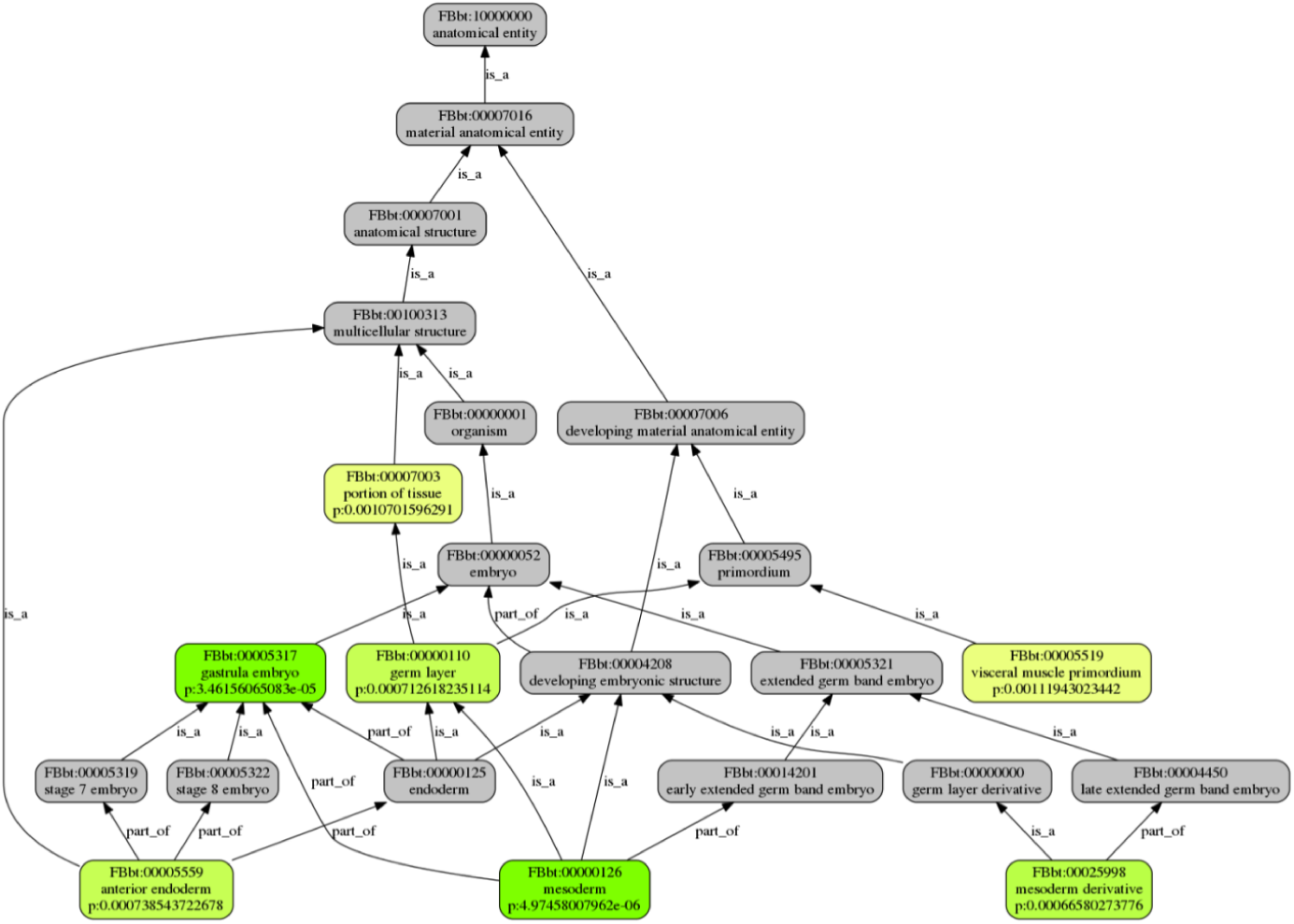
Bio.Ontology enrichment results from [WF10] visualized as a DAG using graphviz.

The fact that our software is implemented to be compatible with biopython and the Galaxy servers allows it to be easily integrated in larger analysis pipelines. In our experience, it makes for an easy integration with differential gene expression analysis in Galaxy platform or analysis of genes targeted by a certain Transcription Factor found in biopython.

## 3 Methods

Let us first define the standard term-for-term enrichment for gene sets. Essentially, we want to find out whether given term in ontology is connected to set of genes representing a biological phenomenon such as differential expression in an experiment.

Let *n* and *m* be the size of the study and population sets and let the *n_t_* and *m_t_* be the number of genes associated with term *t* in those sets. We calculate the probability *P* of drawing *n_t_* or higher number of genes annotated to the term *t* given the distribution (hypergeometric) of the population:

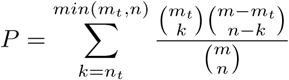

The method described above does not take the dependencies between the terms into account, which effects in some terms being over-represented. One of the solutions to this problem is *parent-child* [GBRV07] method. It calculates the probability described above but given the distribution of genes annotated to the parents of term *t*. Set of genes annotated to the parents of *t* can be computed in two different ways: either as intersection or sum of sets of genes annotated to the each parent. Let *m_p(*t*)_* and *n_p(*t*)_* be the number of genes annotated to the parents from population and study set genes respectively. We calculate the p-value of *t* as follows:

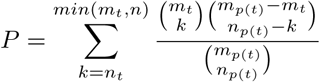

As mentioned before toolbox contains also methods for computing enrichment for gene rankings. Genes are usually ranked based on some biologically relevant criteria (e.g. differential expression between samples) and the goal is to detect any ontology terms enriched in certain extreme parts of the ranking.

First implemented method is *GSEA* applied to ontology terms. Let *S_t_* be the set of genes connected to term *t*. We want to find out whether members of *S_t_* are evenly distributed throughout the rank *L* or not. Sets that contribute to distinction should have their members concentrated at one part of the ranking. First step in *GSEA* method is computing the Enrichment Score - ES. It is calculated by walking down the *L*, increasing the ES when we encounter gene in *S_t_* and decreasing it when we encounter genes not in *S_t_*. The amount of increase is proportional to the gene position in ranking. Next step is calculating the significance of ES. We permute the ranking of genes many times and for each permutation we compute ES which generates null distribution. Our empirical p-value is then computed according to this distribution. As we compute the ES for each term in ontology last step is adjusting the estimated significance level for multiple hypothesis testing based on Bonferroni correction or Benjamini-Hochberg’s FDR.

The second method is *ranked-parent-child*. We create gene set *S_k_* from each prefix of ranking. Then for each *S_k_* we look for enriched terms using described above *parent-child* method. Now the term *t* might be significant for many prefixes with certain p-value. We choose the smallest one as term *t* enrichment. In our experience, this method was able to return significant results with efficiency on par with *GSEA* method.

## Authors Contribution

BW and KK have designed the software and written the original article. KK has written the original implementation. JHI has improved the software including optimization for efficiency, implemented the galaxy interface and deployed the demo site. All authors read and agreed upon the final version of the manuscript.

## Acknowledgments

We would like to thank the members of Biopython community, and Iddo Friedberg in particular for constructive discussion regarding potential uses of Bio.Ontology within the biopython toolbox. This work was partially supported by the Polish National Center for Research and Development grant No. ERA-NET-NEURON/10/2013 and the National Science Center grant decision number DEC-2014/12/W/NZ1/00463

## Footnotes

1 http://regulomics.mimuw.edu.pl:12347

